# Dehydration triggers anomalous subdiffusion in biomimetic cell membranes

**DOI:** 10.64898/2026.09.01.748409

**Authors:** Madhurima Chattopadhyay, Taras Sych, Erdinc Sezgin, Lukasz Piatkowski

**Affiliations:** Institute of Physics, Poznan University of Technology, Piotrowo 3, 61-131 Poznan, Poland; Science for Life Laboratory, Department of Women’s and Children’s Health, Karolinska Institutet, 17165, Solna, Sweden

## Abstract

Lipid diffusion plays a central role in shaping the structural organization of cell membranes, maintaining lipid homeostasis, and facilitating cellular transport and signaling. The lateral mobility of phospholipids in membranes depends heavily on their hydration state. Furthermore, the activation energy of diffusion increases in conditions of reduced membrane hydration, suggesting that the underlying diffusion mechanism changes upon dehydration. Using two variants of fluorescence correlation spectroscopy (point FCS and scanning FCS) and two membrane reporters, we demonstrate that mild dehydration of phase-separated biomimetic cell membranes alters the lipid diffusion mechanism, resulting in anomalous subdiffusion rather than free Brownian motion. Importantly, the anomalous diffusion parameter, α, decreases significantly upon the initial reduction of the membrane hydration layer, and the effect is fully reversible upon rehydration. These observations strongly indicate the reversible shift in lipid diffusion mode rather than irreversible membrane damage. We propose that this anomalous subdiffusion is caused by the formation of temporarily immobile lipid pockets in the membrane upon dehydration. These results therefore provide important insights into the mechanism of lipid diffusion in membranes undergoing local and transient dehydration, which is an important intermediate step in various biological processes associated with membrane fusion, such as neurotransmission, fertilization, and viral entry.

**Why it matters:** Many biochemical processes, such as cell fusion, neurotransmission, viral entry, and fertilization, involve local, transient membrane dehydration. Therefore, understanding lipid behavior under perturbed hydration conditions is crucial. In this study, we found that phosphatidylcholine lipids undergo a striking transition from free diffusion under fully hydrated conditions to anomalous subdiffusion upon dehydration, which reverses upon rehydration. Our findings reveal that changes in membrane hydration affect not only the rate of lipid diffusion but also the nature of the diffusion process itself. These findings provide new mechanistic insight into the relationship between membrane interfacial hydration and nanoscale lipid mobility. More broadly, our study establishes hydration-controlled supported lipid bilayers as a well-defined experimental platform for investigating anomalous and obstructed diffusion in membrane systems.

## Introduction

Lateral diffusion of lipids is a fundamental process that drives the continuous reorganization of cell membrane components, thereby modulating membrane structural flexibility. Due to its immense importance in processes such as cell signaling, membrane trafficking, and structural reorganization, lipid dynamics have been widely studied in both cellular and model membrane systems^1,2^. Lipid dynamics can be affected by various physicochemical parameters, such as lipid composition, fluidity, viscosity, the presence of nano-hindrances, and compartmentalization^3^. The diffusion coefficients (D) of lipids and proteins in cells have been reported to be significantly lower than those observed in model systems^4,5^. Furthermore, the lateral diffusion of membrane constituents in cellular systems does not always exhibit free diffusion, but rather time-dependent mean square displacement (MSD), where MSD is proportional to t^α^, with α <1, is often observed^6,7^. Such diffusion processes are termed anomalous subdiffusion, with α being the anomaly parameter. This anomaly in lipid diffusivity is generally believed to result from the plasma membrane’s highly complex microenvironment, which is not well reflected in model systems. Common causes of the anomaly in lipid dynamics include the presence of slow-moving obstacles, impermeable domains, transient immobilization, interactions with the cytoskeleton and compartmentalization, although it is likely that not all possible influencing factors have been identified^8–10^.

To investigate the lateral diffusion of lipids, fluorescence-based optical microscopy techniques, such as fluorescence correlation spectroscopy (FCS), fluorescence recovery after photobleaching (FRAP), and single particle tracking (SPT), are typically employed^11^. FCS and SPT are particularly useful for recognizing and understanding the anomalous subdiffusion of lipids and proteins. In SPT, the fluorescent particle is monitored over time and space, providing detailed information about the spatiotemporal trajectories of the moving particle. In FCS, information about both diffusion (D) and α, can be obtained by continuous monitoring of the fluorescence fluctuations of particles in the observation volume of the microscope, or along a line a few micrometers long, for point FCS and scanning FCS (s-FCS), respectively, followed by modeling the autocorrelation curves appropriately. These techniques can distinguish between Brownian and non-Brownian (e.g. anomalous subdiffusion or super-diffusion) motions in both cellular and model membrane systems^12–14^.

Biomimetic model membrane systems, such as supported lipid bilayers (SLBs), have been widely used to study the anomalous diffusion behavior of lipids and to understand its potential origins^12,15–17^. FCS combined with stimulated emission depletion microscopy (STED-FCS) revealed that lipids in both the (liquid ordered) L_o_ and (liquid disordered) L_d_ phases move freely in phase-separated SLBs on mica. In contrast, in SLBs on glass, lipids exhibit anomalous subdiffusion^18^. Furthermore, introducing heterogeneities in the form of different lipid phases with varying lipid compositions or membrane-associated proteins causes anomalies in lipid lateral diffusion^9,12,19^. Similar results have been obtained from molecular dynamics (MD) simulations. Anomalous subdiffusion of lipid-binding Pleckstrin homology (PH) domains has been reported in phosphatidyl-inositol phosphate (PIP)-containing lipid bilayers^20^. Systematic MD simulation studies on membranes of varying lipid composition have shown that lipid subdiffusion depends on lipid chemistry, phase, and the presence of cholesterol in the membrane^21^. Using Monte Carlo simulations, Nicolau et al. demonstrated that anomalous diffusion may also be caused by interactions with fixed obstacles, rafts, and collisions with picket fence posts^8^. Simulations involving varying numbers and sizes of immobile obstructions and different percolation thresholds showed that hindered diffusion increases with an increase in the number of obstructions and a decrease in their size^22^. Despite extensive research into anomalous diffusion in membranes caused by various compositional and mechanical factors, the effect of physicochemical parameters such as viscosity, temperature, and, in particular, hydration, has remained unexplored. Recent studies show that, of the many physicochemical parameters affecting lipid diffusion, water molecules near the polar head group of lipids play a crucial role in modulating their lateral diffusion^23,24^. Interestingly, many biochemical processes involving the fusion of two lipid membranes, such as cell fusion, neurotransmission and fertilization, involve the expulsion of water molecules, leading to local and transient membrane dehydration^25^. Therefore, given the biological significance of membrane fusion events, it is important to understand the precise diffusion behavior of membrane components under perturbed hydration conditions. Unfortunately, the lack of a methodology for preparing lipid membranes with well-defined hydration conditions has hindered the study of the effects of local dehydration on the mobility of membrane components thus far. In this study, we employed a recently devised strategy of slow and gradual dehydration of SLBs to investigate phospholipid lateral diffusion characteristics in SLBs on a mica substrate under decreased hydration conditions. To this end, we used two variants of the FCS technique - point FCS and scanning FCS (sFCS) - as well as two fluorescent membrane reporters, Atto-633 and Abberior STAR

RED (ASR)-functionalized lipids, to ensure that the results were independent of the fluorophores used. The Atto and Abberior dyes have been proven to be reliable choices for FCS experiments due to their photostability and high quantum yield^18^. The experimental results revealed that the phospholipid diffusion mode changes to anomalous subdiffusion in the absence of bulk water, reversing to free diffusion upon rehydration. We speculate that this anomaly arises from the formation of transient nano-hindrances under lower hydration conditions. This study provides a deeper understanding of the lipid diffusion behavior in membranes with a perturbed hydration state, which is relevant to the biological phenomena involving membrane fusion.

## Materials and methods

### Materials

1,2-Dimyristoleoyl-sn-glycero-3-phosphocholine (14:1 PC), egg yolk sphingomyelin (egg SM), and cholesterol were purchased from Avanti Polar Lipids, Alabaster, AL, USA. 1,2-dioleoyl snglycero-3-phosphoethanolamine labeled with Atto 633 (DOPE-Atto 633), Alexa 488 fluorescent dye, 4-(2-hydroxyethyl)piperazine-1-ethanesulfonic acid (HEPES), sodium chloride (NaCl), and chloroform (HPLC grade) were purchased from Merck KGaA, Darmstadt, Germany. 1,2-dipalmitoyl-sn-glycero-3-phosphoethanolamine (DPPE) conjugated with the Abberior STAR RED fluorescent dye (ASR-PE) was purchased from Abberior GmbH, Göttingen, Germany. All the materials were used without further purification.

### SLB preparation

The vesicle deposition method previously reported, with the necessary adaptations, was used to prepare SLBs on a mica substrate^26^. Chloroform solutions containing 14:1 PC, egg SM, and cholesterol at a molar ratio of 1:1:1, as well as 0.002 mol% of ASR-PE and 0.01 mol% of Atto-DOPE, were prepared separately. The chloroform was then evaporated by blowing nitrogen gas over the solution for at least 50 minutes to ensure complete removal of the solvent. The dried lipid film formed on the bottom of the vial was dissolved in an aqueous buffer solution (10 mM HEPES and 150 mM NaCl buffer solution, pH adjusted to 7.4) to form multilamellar vesicles (MLVs) with a lipid concentration of 10 mM. The MLV suspension was vortexed thoroughly to obtain a homogeneous lipid solution. This solution was then diluted tenfold to achieve a final lipid concentration of 1 mM, and distributed into sterilized glass vials for storage at −20°C. The MLV suspensions were bath sonicated for 10 minutes at maximum power to produce a solution of small unilamellar vesicles (SUVs). Freshly cleaved mica was adhered to a #1.5 coverslip using UV-activated Norland 68 adhesive, and a quarter-cut Eppendorf tube was attached on top of it as a water reservoir using medical-grade silicone. 100 μL of the SUV solution was deposited on the mica, followed by the addition of 2 μL of a 0.1 M CaCl_2_ solution and 400 μL of buffer solution. The sample was incubated for 30 minutes and then rinsed well with 10 ml of buffer solution to remove unburst vesicles.

### Preparation of SLBs with lower hydration conditions

To obtain SLBs with lower hydration, the bulk part of the buffer solution was removed from the top of the SLB using a pipette. The sample was then immediately purged with N_2_ gas at a relative humidity (RH) of over 90%. The RH of the N_2_ gas was monitored and controlled using a homemade humidity control setup^24^, gradually decreasing to 80% and 70% RH before increasing back to 85% RH. The SLBs were equilibrated for approximately 10 minutes at a specific RH% before measurement. Finally, 200 μL of buffer solution was added to the SLB to achieve ‘bulk rehydration’ of the membrane.

### Point and scanning FCS data acquisition and analysis

Fluorescence imaging and FCS measurements were performed on SLBs with varying hydration conditions using an inverted LSM 780 (Carl Zeiss, Jena, Germany) microscope fitted with a 40x 1.2 NA water immersion objective with a correction collar. Atto 633 and ASR fluorescent dyes were excited using a He-Ne laser with a wavelength of 633 nm, while Alexa 488 dye was excited using an argon laser with a wavelength of 488 nm. For both point FCS and s-FCS experiments, the pinhole was set to be 1 Airy unit. These adjustments of the pinhole and objective correction collar were routinely performed before all FCS experiments. The confocal volume and structural parameters were calibrated using a 5 nM aqueous solution of the standard calibration dye, Alexa 488, before the FCS measurements on SLBs.

Point FCS measurements were performed using the FCS module in Zen software (Carl Zeiss, Jena, Germany). Each point FCS measurement was taken over 5-10 seconds, with five repetitions at a particular spot at low laser power. For each hydration level, point FCS measurements were recorded in a minimum of 6 different areas of the sample, providing a minimum total of 30 FCS curves at each hydration state. s-FCS experiments were performed in line scan mode by continuously imaging a 2.8 μm line 100,000 times at the microscope’s maximum scanning speed. The line was imaged with 36 pixels and a pixel dwell time of 5.64 μsec, resulting in a scanning frequency of 1032 Hz and a total measurement time of 96 s. All s-FCS measurements were performed in photon-counting mode. The point FCS correlated data were fitted using FoCusPoint software^27^. The s-FCS data were analyzed using FoCusScan software^28^, in which correlation carpets were generated by reprocessing the fluorescence fluctuation data. This involved averaging a few individual FCS curves (binning 1 or 5 in FoCusScan) and using the local averaging method (10 s) for photobleaching correction. The correlation curves extracted from a single carpet were averaged to produce one or two FCS curves, compensating for the high noise in the s-FCS data, particularly in the case of scarcely hydrated samples. For both point and scanning FCS, the FCS curves were fitted using a 2D free diffusion triplet relaxation model^28^:

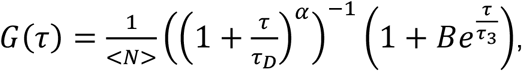

Where < *N* > - average number of particles in the focal volume, *τ*_*D*_ – diffusion time, *α* – anomaly parameter, *τ*_3_ – triplet relaxation time and *B* – amplitude reflecting the triplet population.

The diffusion time for lipids obtained from fitting the FCS data was converted to a diffusion coefficient using the data obtained from calibration according to^29^:

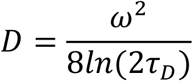

where *τ*_*D*_ is the diffusion time, and ω is the full width at half-maximum of the point spread function. The obtained *τ*_*D*_ of Alexa 488 and its D value in aqueous solution (430 μm^2^/s^30,31^) are used later to calculate the D value of the sample.

Hence:

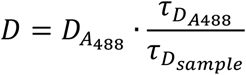

where the diffusion coefficient of the Alexa dye 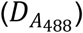 is 430 μm^2^/s, 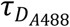 is the diffusion time of the Alexa 488 dye in an aqueous solution, measured routinely during the calibration procedure, and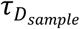 is the diffusion time of the lipids in the sample.

## Results

In this study, we investigated the nature of lipid dynamics in phase-separated SLBs of an equimolar mixture of 14:1 PC, egg SM and cholesterol under varying hydration conditions using point and scanning FCS. Two standard FCS measurement dyes, Atto-633 and ASR, conjugated with the lipid analogs DOPE and DPPE, respectively, were chosen for the experiments to eliminate the possible influence of the fluorophore characteristics on the lipid diffusion. Due to the differences in hydrophobic chain saturation and length, the so-called hydrophobic mismatch, 14:1 PC forms a highly fluid L_d_ phase, while egg SM forms a compact L_o_ phase with a higher cholesterol content (see Figure 1A-D). Atto-633, which is conjugated with the unsaturated lipid DOPE, partitions selectively into the L_d_ phase (Figure 1A-B). Despite being linked to a saturated lipid, ASR-DPPE gravitates towards the fluid, less constricted L_d_ phase due to difficulties encountered by the large fluorophore when entering the compact L_o_ phase^32^ (Figure 1C-D). As the mobility in the L_o_ phase drops to extremely low values after dehydration^23^, hindering the accurate determination of the diffusion coefficient using FCS, this study investigated lipid diffusion specifically in the L_d_ phase.

**Figure 1.**
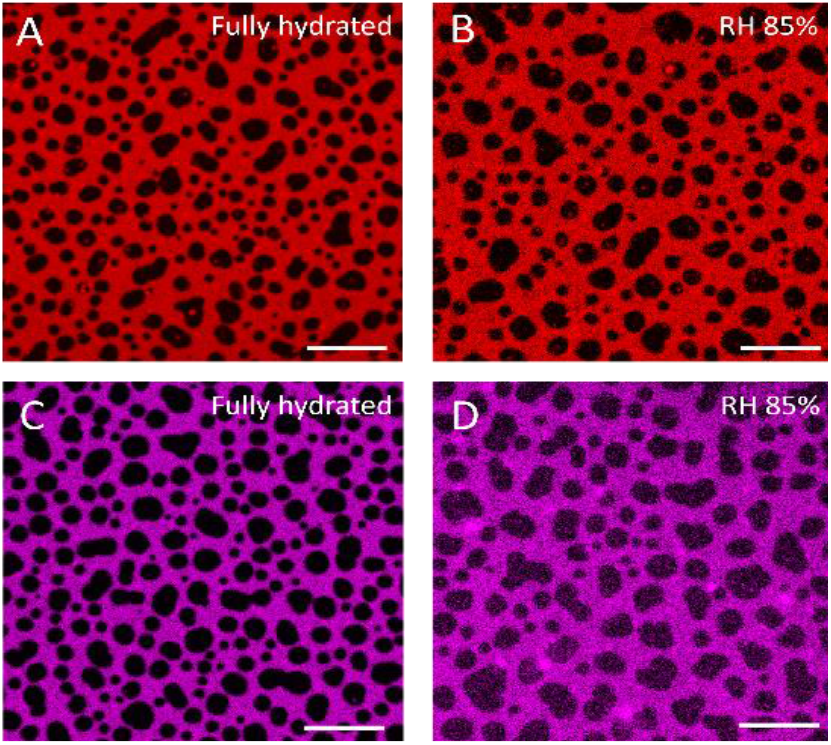
Fluorescence confocal images of a phase-separated SLB containing Atto 633 (A-B) and ASR (C-D)-functionalized membrane reporters. The dark patches correspond to L_o_ domains, while the red and magenta areas correspond to the L_d_ phase labelled with the Atto 633 and ASR dyes, respectively. Scale bar represents 10 µm in all panels.

Point FCS and s-FCS experiments were performed on SLBs in a fully hydrated state and on SLBs that were equilibrated at 90%, 80%, and 70% RH during the dehydration process, and at 85% RH during the rehydration process. Finally, the SLBs underwent bulk rehydration. Removal of the bulk water following our slow and gradual dehydration protocol^23^ had no effect on the SLB structure at the micrometer scale (Figure 1B, D and S1). However, removal of the bulk water led to photobleaching of both the Atto-633 and the ASR dyes; the ASR dye exhibited significantly greater photobleaching than the Atto-633 dye. Reliable point and scanning FCS curves could be obtained for Atto-633 down to 70% RH. Below 70% RH, the extensive photobleaching hindered the reliable analysis of the FCS data (see Figure S2A-B). In contrast, no credible point FCS curves could be obtained for the ASR dye immediately after removing the bulk water, due to significant photobleaching during the experiment (Figure S2C). However, s-FCS is comparatively less sensitive to photobleaching. This enabled us to obtain relatively high-quality FCS curves for ASR-containing membranes down to 70% RH, provided that appropriate photobleaching corrections were applied during analysis. The increased photobleaching of the dyes upon dehydration is likely due to slower lipid diffusion, resulting in a longer residence time within the excitation spot. Furthermore, a change in the photophysical properties of these dyes upon dehydration is suspected, possibly due to a change in polarity of their environment^33^, an effect that requires further investigation. The photostability and high quantum yield of the ASR dye in aqueous solution are well understood, and it is known to be significantly more photostable than the Atto dye^18^. However, given ASR’s highly hydrophilic nature, it is understandable that its photostability is more sensitive to changes in hydration than that of moderately hydrophilic Atto-633.

### Point FCS measurements

FCS is based on temporal fluctuations in fluorescence at the observation spot/volume of the microscope, which are caused by the movement of the fluorophores attached to the target molecules (Figure 2A). Point FCS measurements were performed on six, phase-separated SLBs doped with Atto-633-DOPE. Three of the samples were measured at all the aforementioned RH levels, while the other three samples were only measured at fully hydrated conditions and at 90% or 80% RH. Figure 2B shows the non-averaged, normalized FCS curves for a single SLB containing Atto-633-DOPE in a fully hydrated state (red), at 90% RH (blue), at 80% RH (green) during dehydration, and finally upon bulk rehydration (purple). The FCS curves shift towards longer diffusion times (to the right) with dehydration and back towards shorter diffusion times (to the left) with rehydration. This reconfirms the reversible, hydration-dependent nature of lipid lateral diffusion, in agreement with our previous work^24^. Interestingly, the diffusion time and the shape of the FCS curve both change upon dehydration, and revert with bulk rehydration. This change in the shape of the autocorrelation curves is the first indication that the lipid diffusion in perturbed hydration conditions differs from that in fully hydrated membranes. Figures 2C and 2D show the fitting of the FCS curves with and without fixing the α parameter at 1. The FCS curves obtained for the fully hydrated SLBs can be well fitted by a single-component free diffusion model (Figure 2C). However, the FCS he SLB after the removal of bulk water could not be described by a single free diffusion model (Figure 2C). Consequently, we fitted the data obtained for SLBs at reduced hydration states with two models: i) single-component anomalous diffusion; and ii) two-component free diffusion. In the first model, the α parameter was released to correct for the potential diffusion anomaly, rather than being kept constant at 1. In the second model, two separate populations of freely diffusing particles were considered, and their fractions were released while α remained constant at 1. Both the free α model (Figure 2D) and the two-component free diffusion model (Figure S3) provided a good mathematical fit to the FCS curves. The next step was therefore to check the physical credibility of the extracted parameters. The D values obtained from the two-component free diffusion model did not fall within the generally accepted range of lipid diffusion times in either the gel or fluid phases^34^. The fast component (∼25% of the total lipids) was found to be seven to ten times faster than the standard lipid diffusion in the L_d_ phase, while the slower component (∼75% of the total lipids) was five to ten times slower. Such a significant increase in the D value of the first component upon dehydration is improbable. Therefore, the possibility of two freely diffusing populations was dismissed. In contrast, the D values obtained by fitting the FCS curves with a floating α parameter were physically plausible and consistent with our previous FRAP results^23,24^. Therefore, a model with a floating alpha parameter was chosen. Upon careful analysis of the FCS curves, a two-fold drop in D values was observed: from 2.7 ± 0.79 μm^2^/s (fully hydrated) to 1.3 ± 0.70 μm^2^/s (70% RH), followed by an increase in D with rehydration (2.97 ± 1.05 μm^2^/s for bulk rehydration) (Figure 2E). Notably, the α parameter was found to be very close to 1 for fully hydrated SLBs (1.05 ± 0.17) (Figure 2F), indicating free Brownian motion. Interestingly, upon removal of bulk water, α decreased to 0.69 ± 0.14 at 90% RH (Figure 2F). Following further dehydration to 70% RH (0.74 ± 0.15) and subsequent rehydration to 85% RH (0.71 ± 0.16), α remained largely constant within the error bar. After bulk rehydration, it returned to 1.03 ± 0.09. This significant yet reversible reduction in α strongly suggests that lipids undergo anomalous subdiffusion in the absence of a bulk water layer. To eliminate fluorophore bias in these experiments, the same point FCS measurements were attempted using an ASR-DPPE fluorescent probe. However, reliable data could not be obtained due to significant photobleaching of the fluorophore, as discussed previously.

**2.**
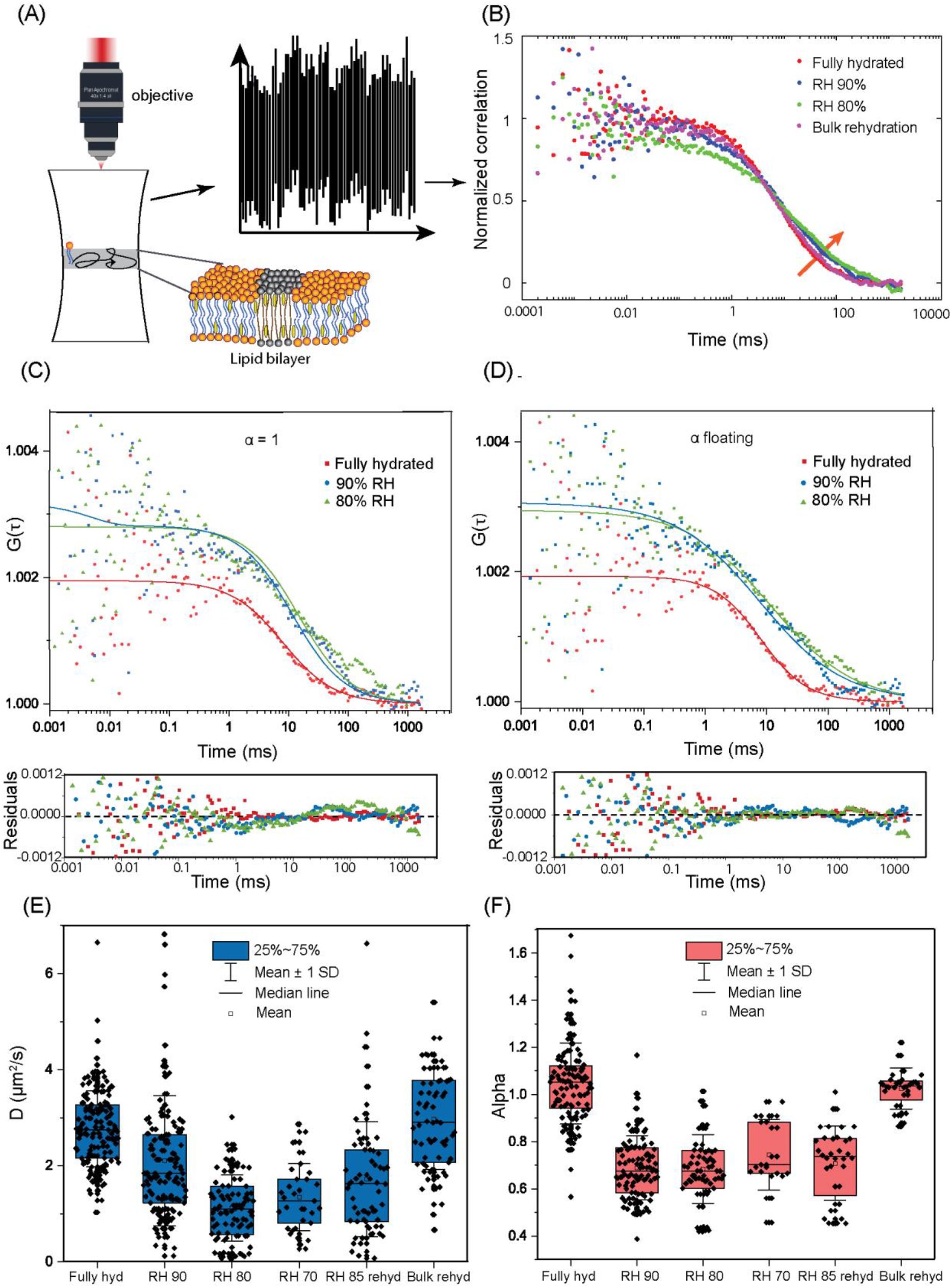
(A) Schematic representation of the point FCS experiment on an SLB. (B) Normalized autocorrelation curves obtained from point FCS measurements on SLBs doped with Atto-633-DOPE in a fully hydrated state (red), at RH 90% (blue), RH 80% (green), and upon bulk rehydration (magenta). The orange arrow indicates the shift and change in shape of the FCS curves upon dehydration. (C)-(D) FCS autocorrelation curves and fitting residuals for SLBs in a fully hydrated (red) state and after equilibration to RH 90% (blue) and RH 80% (green) fitted with α=1 (C) and with floating α values (D), respectively. D (E) and α (F) values for L_d_ phase lipids at varying hydration conditions by fitting with floating α values.

### Scanning FCS measurements

Scanning FCS has previously been used to distinguish between free and hindered diffusion^35^. Rather than recording fluorescence fluctuations at a single point, scanning FCS involves systematically collecting the fluorescence intensity along a selected line multiple time. This generates an autocorrelation carpet comprising several autocorrelation curves that correlate fluorescence fluctuations at each pixel of the line (Figure 3A-B). The diffusion time is then extracted by fitting these curves, in a manner similar to that used for point FCS data. This provides insights into the spatial heterogeneity and a significant statistical averaging of the mobility while compromising little on the temporal resolution. Crucially, in the context of these particular experiments, the effect of photobleaching is less pronounced since a single observation spot is not continuously illuminated by an excitation laser in scanning FCS. Therefore, to validate our point FCS findings, s-FCS measurements were performed on SLBs containing Atto-633-DOPE and ASR-DPPE. Thanks to ASR-DPPE’s comparatively lower photobleaching in s-FCS, reliable correlation data could be obtained in this case. Furthermore, to increase the signal-to-noise ratio, the correlation traces from a single correlation carpet were averaged to yield one or two curves for the samples with lower hydration. Although the averaging leads to a loss of the spatial information about the diffusion, this was deemed acceptable given that no spatial heterogeneity was identified, as evident from Figure S4. Figure 3B presents representative correlation carpets for a fully hydrated SLB and an SLB equilibrated to 90% RH containing ASR-DPPE dye. The averaged and normalized autocorrelation curves for L_d_ lipids in an SLB with ASR-DPPE dye at fully hydrated, 90% RH and 80% RH conditions are shown in Figure 3C. The FCS curves shift to the right as RH% decreases, indicating a reduction in the diffusion coefficient value. Furthermore, upon SLB dehydration, a clear change in the shape of the correlation curves is observed, similar to the point FCS data. As with the point FCS data, the correlation curves were fitted using both a floating α model (effective for anomalous diffusion) and a two-component free diffusion model. However, fitting the two-component free diffusion model was not used for the same reason as for the point FCS data. Figure 3D and 3E show the same correlation curves fitted with α = 1 and α floating, respectively. Similarly, fitting of three representative autocorrelation curves for SLBs containing an Atto-633-DOPE probe, each averaged over a single carpet, with α = 1 and α floating, is presented in Figure 3F-G. The fits and their residuals show that the free diffusion model (with α = 1) fits the experimental data well in the fully hydrated state but not at RH levels of 90% and 80%. Conversely, allowing α to vary significantly improves the quality of the fit at lower hydration levels. Thus, the FCS curve fitting for both dyes confirms the occurrence of anomalous subdiffusion in membranes with perturbed hydration. Figures 4A-B and 4C-D show the diffusion coefficient and α values for the Atto-633-DOPE and ASR-DPPE dyes, respectively, as a function of the SLB hydration state. The absolute D values are lower than those obtained from point FCS measurements, but follow the same dehydration and rehydration trend as the point FCS results, irrespective of the dye used. Furthermore, under fully hydrated conditions, α was also found to be approximately 1 (0.97 ± 0.13 for Atto633 and 1.05 ± 0.09 for ASR), decreasing to 0.81 ± 0.11 for Atto633 and 0.85 ± 0.09 for ASR under lower hydration conditions (RH 80%). Again, upon rehydration, α went back up to 0.91 ± 0.09 for Atto and 1.03 ± 0.08 for ASR.

**Figure 3.**
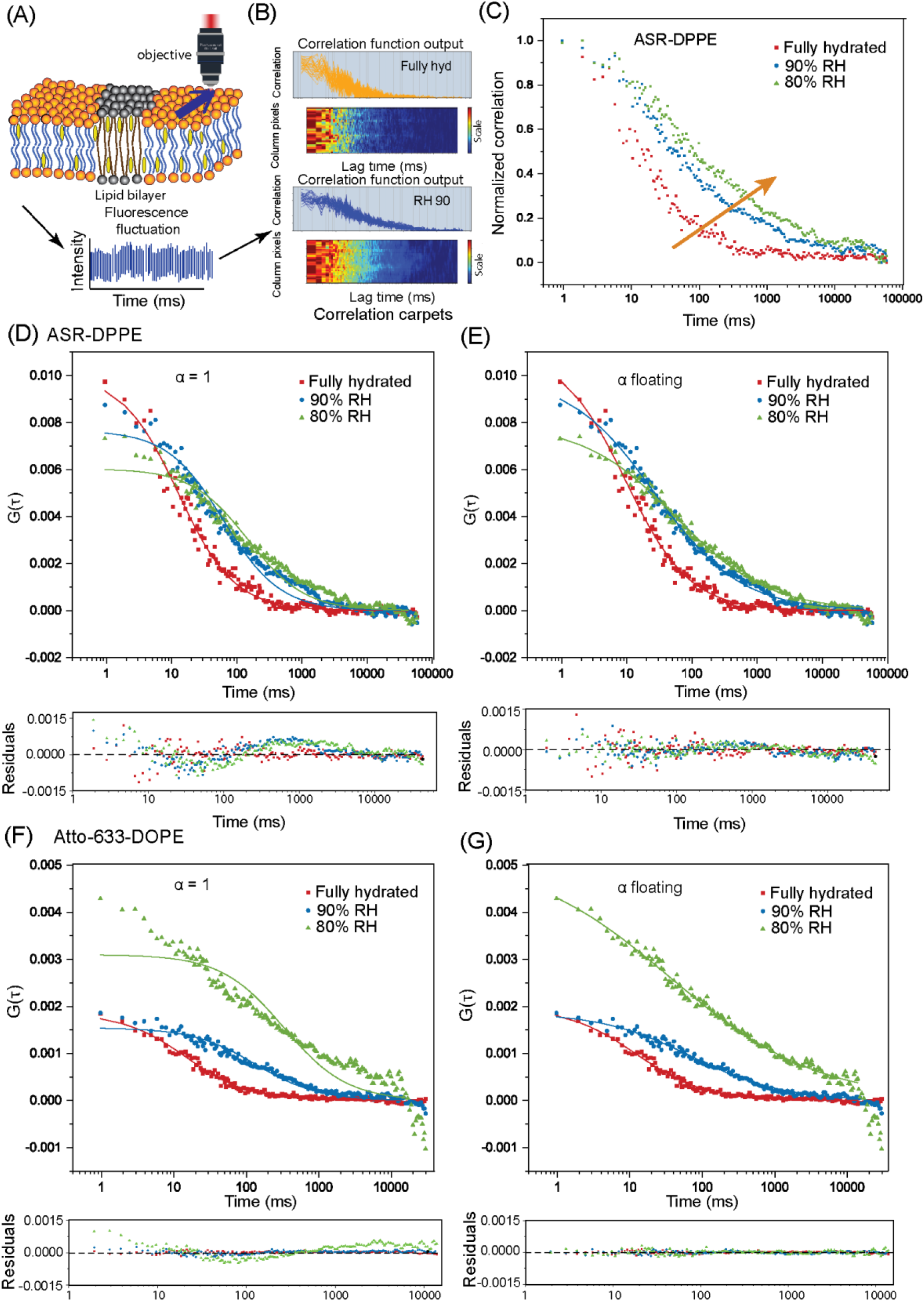
(A)-(C) Schematic representation of scanning FCS on SLB; (B) Representative correlation carpets for SLBs doped with ASR-DPPE in a fully hydrated state and at 90% RH; (C) Normalized correlation curves, each averaged from single correlation carpets in a fully hydrated state (red), at 90% RH (blue) and at 80% RH (green). The orange arrow indicates the shift of the FCS curves upon dehydration. (D-E) s-FCS curves for SLBs with ASR-DPPE at fully hydrated conditions (red) and equilibrated to RH 90% (blue) and RH 80% (green) fitted with α=1 (D) and with floating values of α (E); (F-G) s-FCS curves for SLBs with Atto-633-DOPE at fully hydrated conditions (red) and equilibrated to 90% RH (blue) and 80% RH (green), fitted with α=1 (F) and with floating values of α (G).

**Figure 4.**
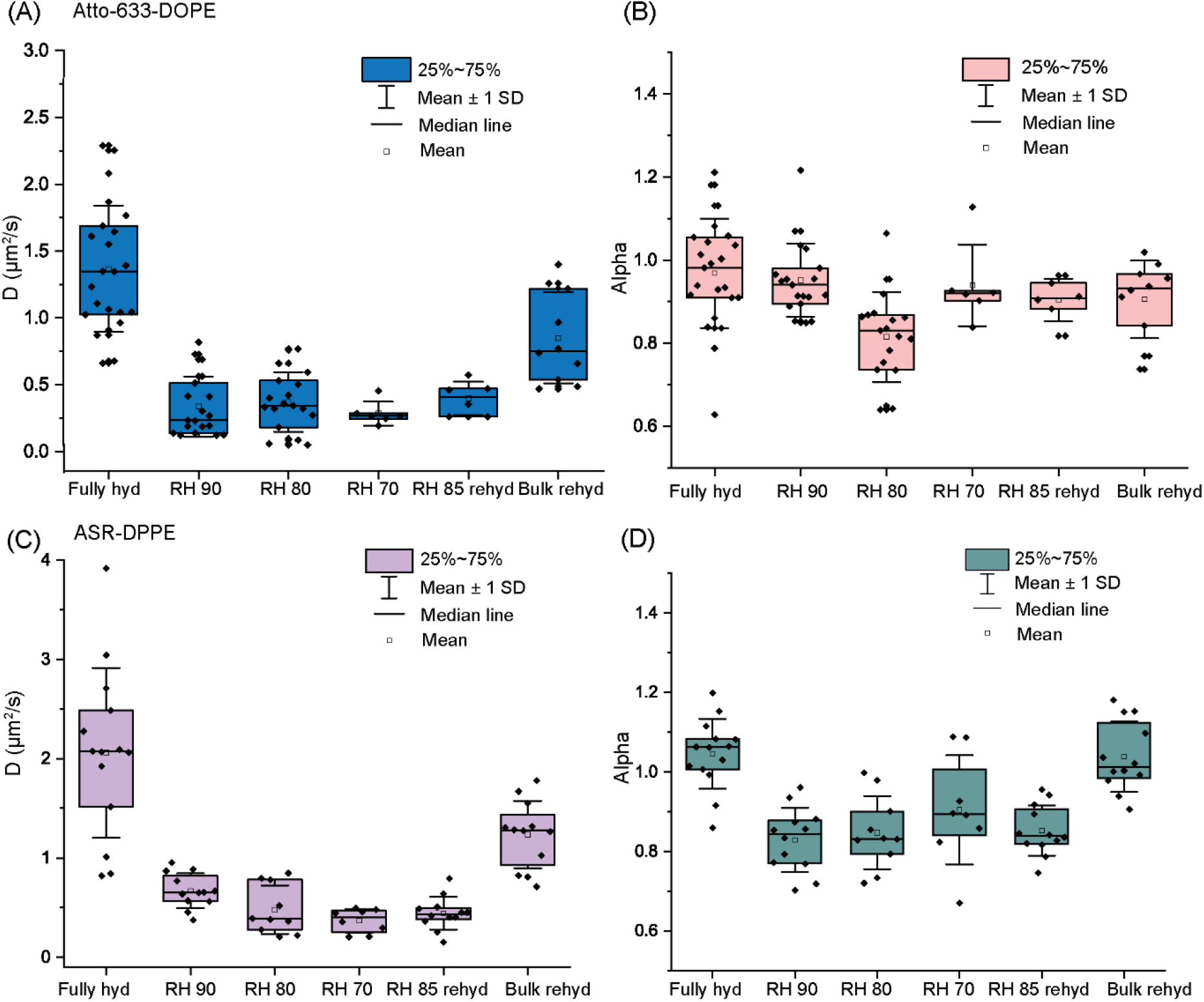
D (A) and α (B) values obtained from s-FCS measurements for L_d_ phase lipids in SLBs with Atto-633-DOPE at varying hydration conditions. D (C) and α (D) values obtained from s-FCS measurements for L_d_ phase lipids in SLBs with ASR-DPPE at varying hydration conditions.

## Discussion

The similar trend in D values acquired from point and s-FCS measurements for different SLB hydration states is consistent with the previously observed trend of hydration-dependent lateral diffusion in FRAP measurements^23,24,36^. Both point FCS and s-FCS data confirm that in the presence of bulk water atop the SLB deposited on the mica substrate, L_d_ phase lipids undergo free diffusion. Conversely, as soon as the direct hydration layer of the membrane is perturbed, the anomaly parameter α drops significantly, reflecting a change in the diffusion mode of PC lipids from free diffusion to anomalous subdiffusion. Notably, the changes in the α parameter are completely reversible: that is, α returns to a value of 1 upon bulk rehydration. The strong reproducibility and reversibility of the de(re)hydration results for both point and s-FCS, as well as for the two benchmark dyes (Atto-633 and ASR), corroborate the idea that the lower α values accurately reflect the diffusion properties of the lipids in SLBs with reduced hydration. Additionally, our previous experiments reported that the activation energy of diffusion increases with dehydration, strongly indicating a change in the diffusion mechanism^24^. This further validates the results of anomalous subdiffusion.

Anomalous subdiffusion is typically observed in the presence of mobile or immobile obstructions, such as traps or domains^9^. When water is removed, the emergence of nanoholes in the membrane that could prevent lipids from diffusing freely would be the first place to look for the cause of the anomalous diffusion behavior of the lipids. However, an atomic force microscopy study revealed that no nanoscopic holes emerge in the L_d_ phase during dehydration^24,37^, which rules out this hypothesis. Therefore, another membrane feature must transiently hinder lipid movement. Our previous study using FRAP demonstrated that the mobile fraction of lipids decreases with dehydration; that is to say, at a given time, a greater number of lipid molecules become transiently immobile^24^. As dehydration occurs, the clathrate-like shell structure around the phosphocholine group breaks apart and disintegrates. As more and more lipids lose their water layer, electrostatic repulsion between the lipid headgroups increases, raising the energy required for diffusion. This could lead to the transient or permanent immobilization of these lipids, which could act as obstructions and cause an anomalous diffusivity pattern. In reality, the water clathrate cage around the lipid headgroups should not be viewed as a concrete, discrete cage structure; rather, the water molecules form a fluid, shell-like layer around the headgroups. Upon dehydration, the perturbation passes on from one lipid to another via the exchange of water molecules in their hydration shells. Therefore, it is reasonable to conclude that their diffusion pattern differs significantly from that of free diffusion.

Previous studies of the phase transition temperature, the height mismatch between the L_o_ and L_d_ phases, and the fluorescence spectra of the membrane-embedded environmental probes suggest that a transition occurs from a disordered liquid phase to a quasi-gel phase upon membrane dehydration^33,37,38^. If the fluid lipids were to transform gradually into the gel phase, the two-component free diffusion model, with D values matching those of the fluid and gel phases, would describe the diffusion data. According to this model, there would be a gradual increase in the fraction of lipids with gel-phase-like diffusion coefficient. However, the experimental data presented here clearly show that this is not the case. Instead, single-component anomalous subdiffusion was found to describe the lipid dynamics after dehydration better. The anomaly parameter α decreases to approximately 0.8 upon removal of bulk water and remains constant thereafter, despite a steady decrease in the diffusion coefficient as hydration decreases. Assuming that, upon dehydration, an increasing number of lipids experience the electrostatic repulsion from the head groups acting as obstacles, α could gradually decrease as the mobile fraction decreases. Generally, the anomaly parameter decreases with increasing obstacle concentration until the percolation threshold for the anomaly is reached, at which point diffusion becomes anomalous at all timescales^39^. A constant α therefore suggests that, as soon as bulk water is removed, the number of obstacles (i.e., transiently immobile lipid pockets) reaches the percolation threshold of anomalous subdiffusion. Accordingly, with further dehydration, even if the number of immobilized lipid pockets increases, α does not decrease further. Previous studies have shown that dehydration leads to a dynamic redistribution of cholesterol in phase-separated membranes, whereby cholesterol migrates from the liquid-ordered phase to the liquid-disordered phase upon dehydration^40^. The increased cholesterol content in the L_d_ phase leads to a reduction in the diffusion coefficient of lipids in this phase. However, a decrease in hydration from bulk hydration to approximately 70–80% RH would increase the cholesterol fraction from approximately 30% to 40%. This, in turn, results in only an approximately 10% decrease in the diffusion coefficient of L_d_ lipids^41^. Clearly, dehydration-induced remodeling of the lipid membrane composition alone cannot account for the observed effects or their magnitude. When investigating the cause of the anomalous subdiffusion of scarcely hydrated lipids, it is also important to consider the possibility of altered interactions between the lipids and the mica substrate under reduced hydration conditions. Although a thin layer of water is presumed to exist between the lower leaflet and the mica substrate, the interactions between the lipids in the lower leaflet and the substrate are not well understood. Honigman et al. demonstrated the free diffusion of phosphatidylcholine (PC) lipids on mica under fully hydrated conditions. In contrast, the same lipids undergo anomalous subdiffusion on an acid-cleaned glass surface^34^. The data presented here confirm that, under fully hydrated conditions, the diffusion is free on mica. At lower hydration levels, however, the interaction between the PC lipids and the mica surface may change. This would result in more prominent anomalous diffusion of the lipids in the lower leaflet, as well as a different diffusion coefficient for these lipids. Nevertheless, a previous study of lipid diffusion in a lipid bilayer with a fluorescence-quenched upper leaflet revealed that the mobility of the lower leaflet mirrored that of the entire bilayer during dehydration^24^. This suggests that the anomaly in lipid diffusion is not, at least solely, an effect of changes in the lipid-mica interactions upon dehydration.

Further insights into the nature of the observed anomalous subdiffusion could be obtained by performing spot variation FCS or STED-FCS, which allows various types of anomalous subdiffusion to be discerned, such as hop diffusion or transient immobilization^42–44^. However, STED-FCS experiments on a dehydrated system are extremely difficult to perform due to the low photostability of the chosen dyes under reduced hydration conditions. STED-FCS uses very high laser powers, resulting in strong photobleaching of ASR or Atto-633 dyes. Therefore, the search for a photostable dye that can be used in low hydration conditions and when exposed to high laser power continues.

## Conclusions

Using point FCS and s-FCS measurements with two benchmark fluorescent dyes, we demonstrated that PC lipids in SLBs on a mica substrate undergo anomalous subdiffusion at reduced hydration conditions. This contrasts with the free diffusion observed for the same system at fully hydrated conditions. We hypothesize that this anomaly in lipid diffusion is caused by the dehydration-induced transiently immobile lipid pockets acting as nanohindrances to moving lipids and resulting in an obstructed diffusion pattern. Transient dehydration is an important intermediate step in many biological phenomena, including cell fusion, neurotransmission, fertilization, and viral entry. Therefore, a detailed understanding of the lipid dynamics at a molecular level in dehydrated conditions is crucial for unraveling the mechanisms underlying such biological events. This study provides new insights into the lipid diffusion patterns of PC lipids in dehydrated membranes. Furthermore, it advances the field of in vitro research and the quest to determine the various causes of anomalous diffusion in biological systems. Further research using suitable fluorescent dyes and employing spot-variation FCS/STED-FCS on these systems may provide additional information about the precise nature of the observed anomaly in the diffusion of scarcely hydrated lipids.

## ASSOCIATED CONTENT

### Supporting material description

Supplemental figures S1-S4 (PDF)

### Data availability

The data underlying this study are available from the corresponding authors.

### Author contributions

MC: Conceptualization, Methodology, Investigation, Formal analysis, Visualization, Writing-Original draft preparation, Funding acquisition. TS: Methodology, Software, Validation, Writing-Reviewing and Editing. ES: Supervision, Resources, Writing-Reviewing and Editing. LP: Supervision, Project administration, Resources, Funding acquisition, Writing - Review & Editing. The manuscript was written through contributions of all authors. All authors have given approval to the final version of the manuscript.

### Declaration of interests

The authors declare no competing financial interests.

## Acknowledgments

MC acknowledges the financial support from EMBO Scientific Exchange grant 9439. ES has been supported by Swedish Research Council Grants (grant no. 2020-02682, 2024-02993, 2024-00289 and VR-RFI 2016-00968), Wellcome Leap’s Dynamic Resilience Program (jointly funded by Temasek Trust), CancerFonden (25 4592 Pj) Karolinska Institutet (2026-02401; 2024-03250; 2024-03341; 2022-00803), Cancer Research KI (2024-03488), Strategic Research Programme in Diabetes at Karolinska Institutet, Human Frontier Science Program (RGP0025/2022), Longevity Impetus Grant from Norn Group, Hevolution Foundation and Rosenkranz Foundation. ES is an EMBO Young Investigator (YIP-2025). LP has been supported by the National Science Centre under the OPUS grant (2020/37/B/ST4/01785) and SONATA BIS grant (2022/46/E/ST4/00132). We thank the SciLifeLab Advanced Light Microscopy facility and National Microscopy Infrastructure Sweden for their support on imaging.

## Table of content

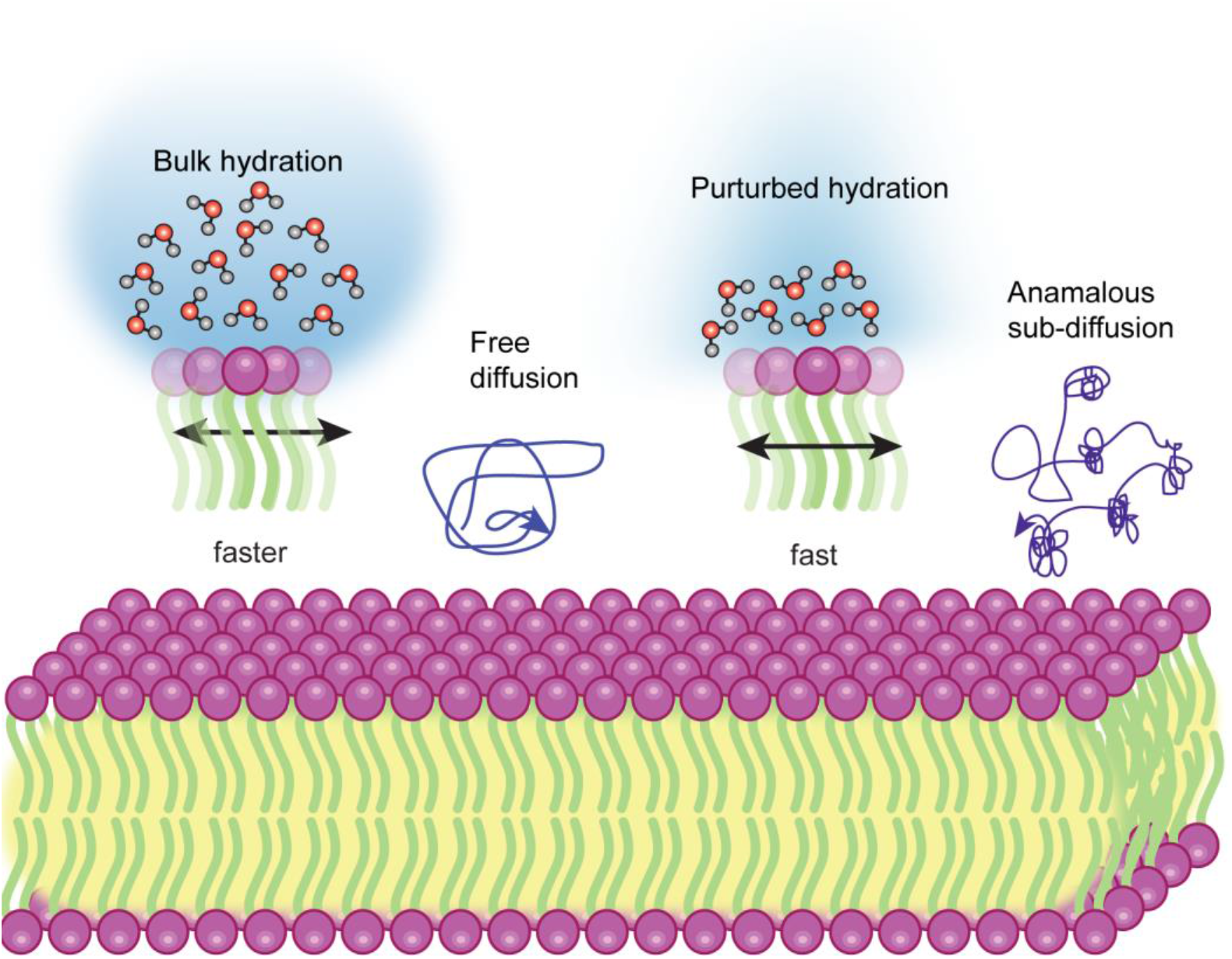

## Notes

### Competing Interest Statement

The authors have declared no competing interest.

