## Supplemental figures for "Dehydration triggers anomalous subdiffusion in biomimetic cell membranes"

For

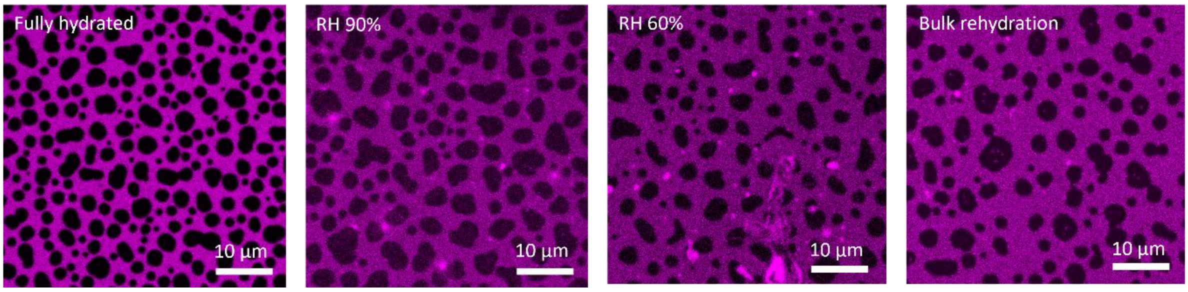


Figure S1: Confocal images of SLBs at various levels of hydration.

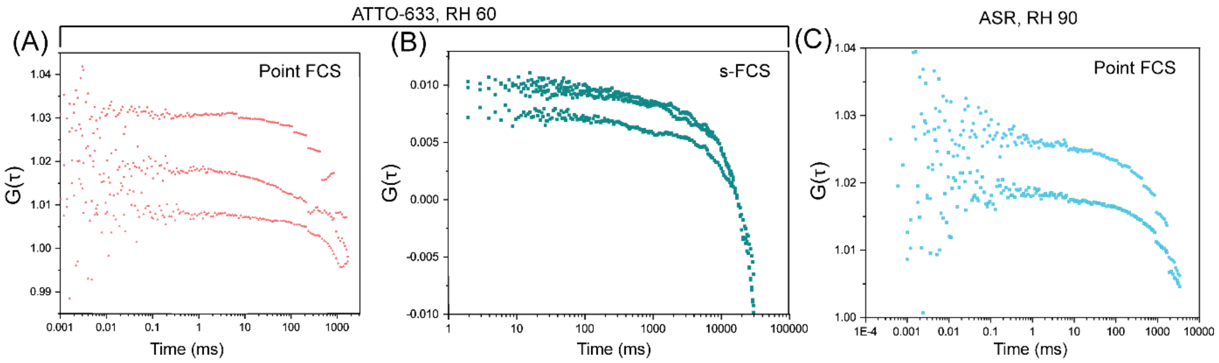


Figure S2: Representative point FCS (A) and s-FCS (B) curves for SLBs with Atto-633-DOPE at lower hydration conditions (<70% RH). Extensive photobleaching of Atto-633 hindered the acquisition of high-quality FCS curves below 70% RH. Photobleaching of the ASR dye occurred shortly after removal of bulk water, preventing acquisition of reliable point FCS curves.

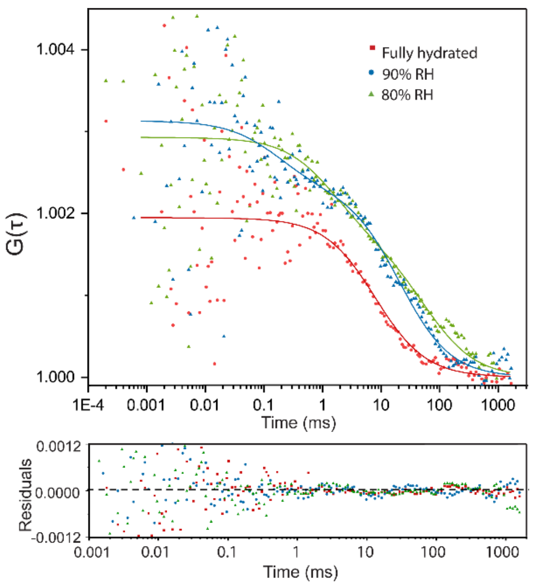


Figure S3. Results of the fitting with the two-component free diffusion model, alongside the residuals of representative point FCS curves for lipids in SLBs under fully hydrated conditions, at 90% RH, and at 80% RH.

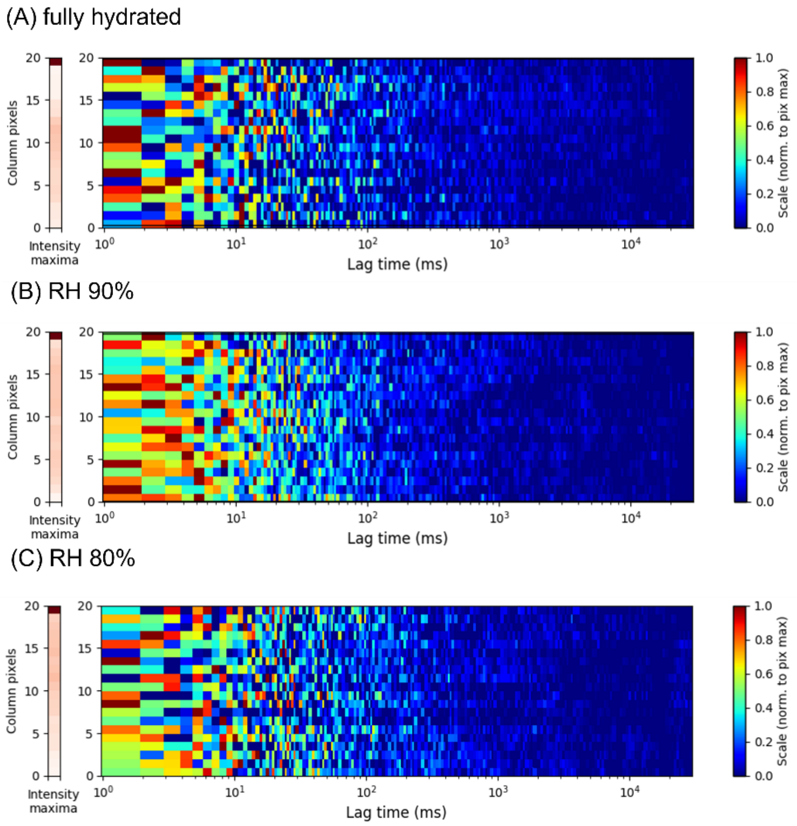


Figure S4. Representative sFCS correlation carpets without averaging: (A) fully hydrated condition; (B) 90% RH; (C) 80% RH. No significant spatial heterogeneity is observed upon dehydration.
